# Individual-specific strategies inform category learning

**DOI:** 10.1101/2024.09.26.615062

**Authors:** Jared S. Collina, Gozde Erdil, Mingyi Xia, Christopher F. Angeloni, Katherine C. Wood, Janaki Sheth, Konrad P. Kording, Yale E. Cohen, Maria N. Geffen

## Abstract

Categorization is an essential task for sensory perception. Individuals learn category labels using a variety of strategies to ensure that sensory signals, such as sounds or images, can be assigned to proper categories. Categories are often learned on the basis of extreme examples, and the boundary between categories can differ among individuals. The trajectories for learning also differ among individuals, as different individuals rely on different strategies, such as repeating or alternating choices. However, little is understood about the relationship between individual learning trajectories and learned categorization. To study this relationship, we trained mice to categorize auditory stimuli into two categories using a two-alternative forced choice task. Because the mice took several weeks to learn the task, we were able to quantify the time course of individual strategies and how they relate to how mice categorize stimuli around the categorization boundary. Different mice exhibited different trajectories while learning the task. Mice displayed preferences for a specific category, manifested by a choice bias in their responses, but this bias drifted with learning. We found that this drift in choice bias correlated with variability in the category boundary for sounds with ambiguous category membership. Next, we asked how stimulus-independent, individual-specific strategies informed learning. We found that the tendency to repeat choices, which is a form of perseveration, contributed to long-term learning. These results indicate that long-term trends in individual strategies during category learning affect learned category boundaries.

## Introduction

Categorization is a basic cognitive task that is fundamental to perception and everyday behavior^1–3^. Individuals apply discrete category labels to incoming sensory inputs, including images, sounds or perceived movements, to understand and respond to our environment. Distinguishing pictures of cats from those of dogs or recognizing faces of specific individuals are standard examples of visual categorization and are fundamental to visual cognition. Categorization is also essential to auditory communication, be it distinguishing between vocalizations of different species, or assigning speech sounds to different syllables or words. In some situations, auditory category labels are learned through exposure and practice^4^, for example when learning a new language as an adult.

The process of category learning can take a long time and differ in the behavioral strategies used between different individuals^5^. Initial biases (such as favoring /r/ sounds over /l/ sounds) and a variety of potential strategies (such as repeatedly sampling information from one category and ignoring a second category as opposed to alternating between the two categories) result in a range of learning trajectories. These trajectories shape the learning outcome, for example categorization accuracy^6^. However, much of the research on categorization focuses on the behavior of subjects that have already learned the task, rarely quantifying the extent to which that behavior is informed by individual differences in category learning.

Categorization of novel or category-ambiguous stimuli is informed by the way that the category information was acquired^7^. Throughout category learning, choices of category labels become more correlated with the category of the stimulus. At the same time, individuals exhibit individual biases in the choice, preferring one category over another^8^. These biases can drift in the process of learning, even in trained subjects. This reflects changes in categorization strategy^9^, and can be linked to an exploration/exploitation trade-off^10,11^. However, the quantitative relationship between individual learning trajectories and learned categorization is not well-understood. Here, we focus on the link between drift in choice bias during category learning and the variability in category boundaries. Specifically, we hypothesized that inter-subject variability of choice bias during learning correlates with category-boundary consistency across testing sessions. Analyzing how individual strategies inform performance outcomes will enrich our understanding of the underlying mechanisms of category learning.

To establish a quantitative relationship between individual strategies and outcomes of category learning, we trained mice to categorize auditory stimuli and to report their choices while performing a two-alternative forced choice (2AFC) task. Mice were trained on extreme high and low tone frequencies and tested on intermediate tone frequencies. Using a machine-learning approach, we extracted the stimulus-independent components of the decision-making process during learning and studied their relationship with categorization parameters after learning. We found that the drift in the choice bias during training is predictive of the stability of the category boundary across sessions when we presented mice with stimuli from ambiguous categories. We additionally hypothesized that stimulus-independent strategies serve as major factors contributing to long-term learning. We found that variability in the drift of the trajectories during early learning could be captured by a model that allowed for perseverative behavior. These results demonstrate that individual stimulus-independent components of learning trajectories are an essential component of category learning, relate to learning outcomes, and should be considered in future categorization studies.

## Results

We trained mice to categorize auditory stimuli and to report their category choices with a 2AFC task^12^. In early parts of training, mice categorized tone bursts that were uniformly drawn from either a low-frequency distribution (6-10 kHz) or a high-frequency distribution (17-28 kHz). Each of these training stimulus sets consisted of 10 tone bursts with central frequencies spanning the respective frequency range. Mice turned a wheel either clockwise or counterclockwise to indicate the category membership of a stimulus (Figure 1A). On average, it took the mice 6844±673 (N=19) trials to reach our pre-defined testing threshold of 75% accuracy for each category (Figure 1D). During training, mice had a (non-significant) tendency to choose low-category stimuli (average accuracy asymmetry = 9.8%±12.7% toward the low category; N = 19, Figure 1D). Across mice, there was not any consistent initial category-accuracy asymmetry (average initial accuracy asymmetry = 3.2%±30.3% toward the low category stimuli; t = -0.253, p = 0.803, One-sample Student’s t-test, N = 19). Once mice reached the testing stage, we introduced 10 novel tone bursts with frequency values between the two training frequency regions (10-17 kHz) (Figure 1C). These stimuli were presented on 20% of trials. We did not reward behavioral responses to these test stimuli.

**Figure 1.**
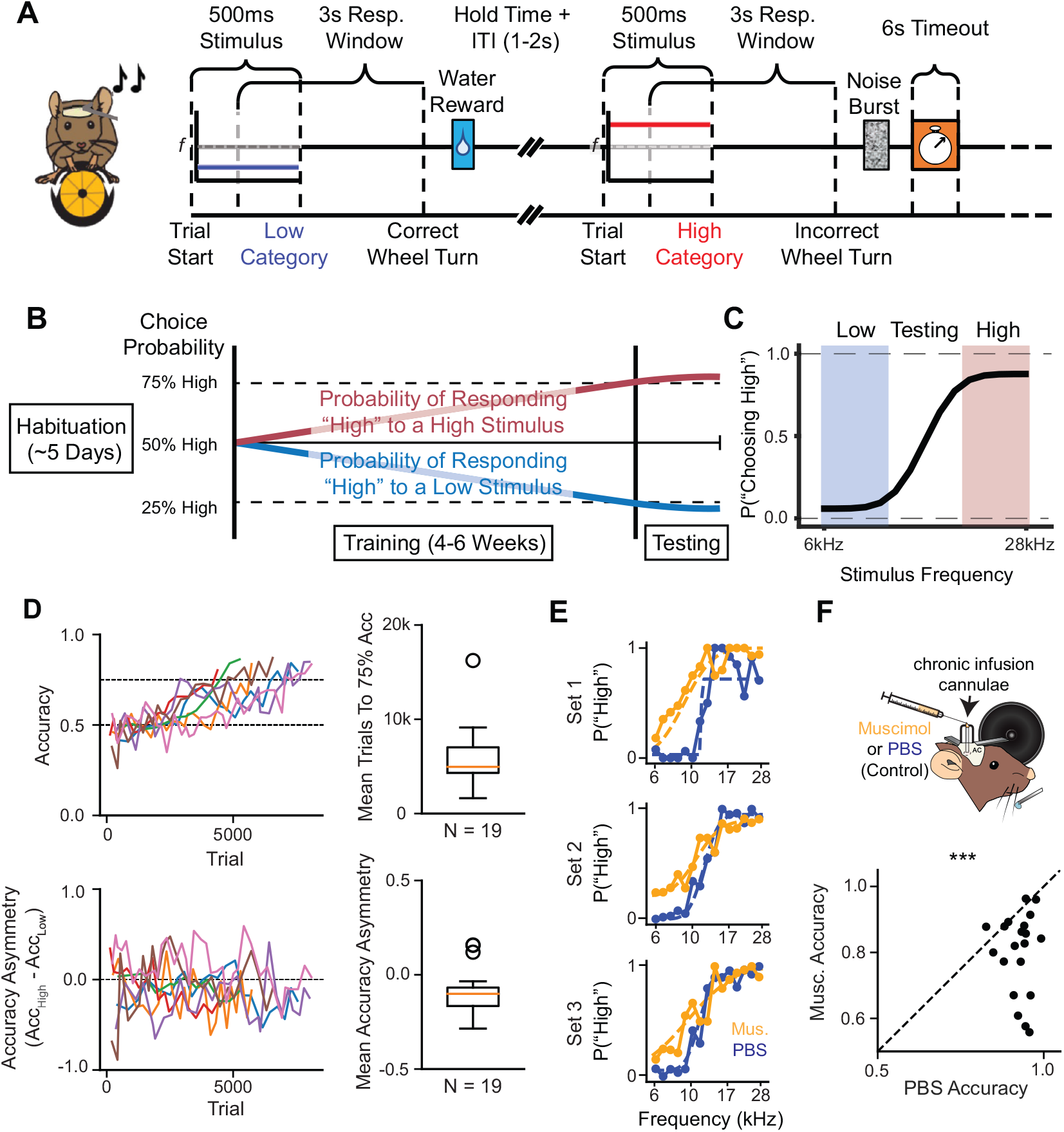
Mouse accuracy on auditory categorization task, which is driven by the auditory cortex, improves over weeks. A, Mice initiated each trial by holding the wheel. We presented a tone burst for 500 ms. After a 100ms delay, the mice were allowed to indicate their choice: a counterclockwise turn indicated that the tone burst was in the “low-frequency” category, whereas a clockwise turn indicated a “high-frequency” category choice. We rewarded the mice on correct trials when the tones were from the low-frequency or high-frequency distribution. Incorrect trials led to a 6 second timeout. For probe tones during the testing stage, the mice were rewarded randomly on a trial-by-trial basis. B, Example learning trajectory. The probability that the mouse turns in the correct direction for high- and low-frequency sounds will increase over time. C, Example psychometric performance on the testing stage. D, Top left: 7 sample mice are shown. Top right: Mice took ∼7000 trials to reach 75% accuracy for each category choice, which was our threshold to start the testing phase with the probe tones. Whiskers represent quartile ranges. Bottom left: Preferred direction (i.e., higher choice accuracy) often switched across sessions. Bottom right: On average, mice tended to have slightly higher accuracy for low-category stimuli compared to high-category stimuli. Whiskers represent quartile ranges. E, Example muscimol and control behavioral data for a single mouse. Each “Set” represents consecutive control and inactivation sessions. F, During the testing stage, we found that bilateral muscimol inactivation of ACx decreased performance accuracy. In all plots: n.s., not significant; *p<0.05; **p<0.01; ***p<0.001; ****p<0.0001.

The auditory cortex (ACx) is known to play a role in auditory perceptual decision-making and learning^13,14^. There is significant evidence that neurons in the auditory cortex encode stimulus category membership^15–18^. There is also evidence that ACx neurons can encode behavioral choice^19,20^. To test whether the ACx is involved in our task, we bilaterally inactivated the ACx of trained, behaving mice using the GABA-A receptor agonist muscimol (N = 20 session comparisons from 6 mice, Fig. 1E, F). We found that when we inactivated the ACx, the mouse performance accuracy for the trained stimulus regions decreased significantly by 11% ±2% (s = 11, p = 0.0002, Wilcoxon sign-rank test), indicating the ACx does play a role in our auditory categorization task.

To study the relationship between the trajectories of the categorization behavior of the mice during learning and categorization of stimuli with ambiguous category memberships, we initially focused on the choice bias in the decision process. In our task, the choice bias was not limited to one direction throughout learning (Fig. 1D). Across all mice, 22.9% ±11.1% (N = 19) of sessions had choice biases in the opposite direction from each mouse’s overall choice bias. This indicates that the choice bias might reflect a dynamic preference that could inform the location of the internal category boundary. We isolated the choice bias using a generalized linear model (GLM) (Figure 2A). This extracted GLM choice bias specifically captures stimulus-independent drift in category preference. We found that mice with more (less) variability in the GLM choice bias during the final training session had more (less) variability in their category boundary during the first five testing sessions (*ρ* = 0.67, p = 0.002, Spearman correlation) (Figure 2D). If the variability in GLM choice bias reflected a generally noisy or inconsistent psychometric curve, then it would be correlated with variability in the other parameters of the standard model as well. However, the drift in the GLM choice bias was not significantly correlated (at the p = 0.05 threshold) with variability in the psychometric slope (*ρ* = 0.44, p = 0.07, Spearman correlation), variability in the psychometric guess rate (*ρ* = 0.057, p = 0.83, Spearman correlation), and variability in the psychometric lapse rate (*ρ* = 0.14, p = 0.57, Spearman correlation) (Figure 2E-G). These results indicate that choice bias during category learning reflects a dynamic preference that mice rely on when drawing category boundaries.

**Figure 2.**
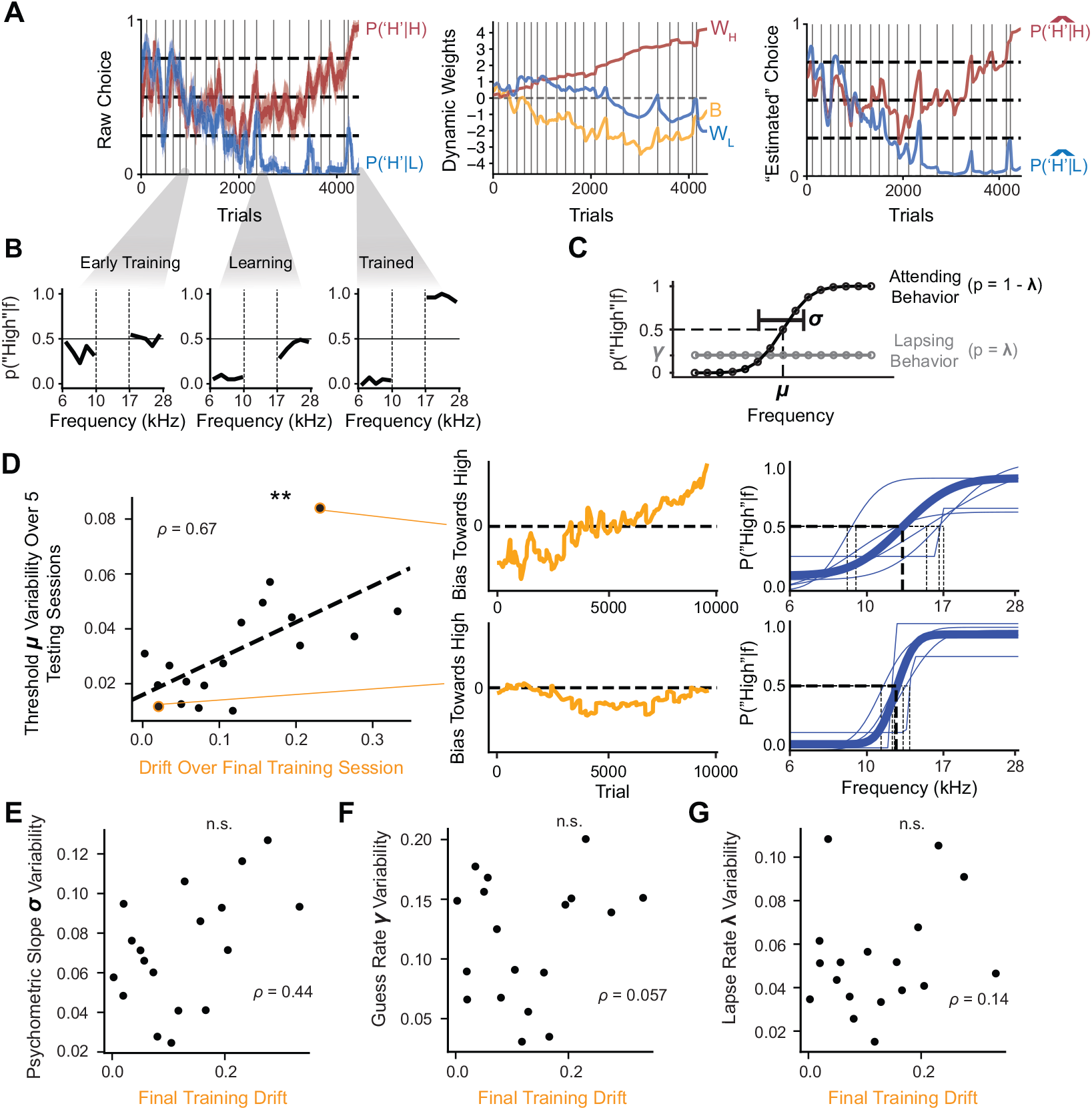
Choice bias during category learning reflects a dynamic preference that mice rely on when drawing category boundaries. A, The PsyTrack model extracts dynamic weights for each category as well as a dynamic choice bias term. Error bars represent 95% CIs. Right: The original traces for the category-conditioned accuracies throughout learning can be estimated from the extracted weights by taking the logistic transformation of the weights. B, The session-level behavioral choice data at three different points throughout learning. C, Visualization of psychometric curve parameters. The animal behavior is modeled as a weighted average of attending behavior (as captured by a sigmoid) and lapsing behavior, which is stimulus-independent. D, Left: The drift over the final day of training is correlated with across-mouse variability in the location of the psychometric threshold during the first 5 testing days. Center: Two examples of isolated GLM choice bias traces. Top panel shows a trace with high variability, bottom panel shows a trace with low variability. Right: Psychometric curves from the testing stage for the same mice, demonstrating that a mouse with higher (lower) variability in dynamic weight has higher (lower) variability in threshold location. Thin lines are individual sessions, thick line is generated from average parameters across the first five sessions. E, The drift over the final day of training is not significantly correlated with the across-mouse variability in the psychometric slope. F, The drift over the final day of training is not significantly correlated with the across-mouse variability in the psychometric guess rate. G, The drift over the final day of training is not significantly correlated with the across-mouse variability in the psychometric lapse rate. In all plots: n.s., not significant; *p<0.05; **p<0.01; ***p<0.001; ****p<0.0001.

We found that the variability in the GLM choice bias at the end of training was predictive of category boundary stability across sessions. Therefore, we focused on extracting patterns in choice bias during learning to understand what drives inter-subject variability in the decision-making process. We generated trajectories for each mouse’s choice bias across learning and applied dynamic time-warping (DTW) clustering to identify the two most prominent clusters (Figure 3A). We found that the key aspect that differentiates the two clusters is whether the initial choice bias persists throughout learning or if the choice bias drifts towards the initially unpreferred category during the early stage of learning (Figure 3B). The distinction between “stationary” and “drifting” trajectories, respectively, is reflected in the trajectories for the response probabilities to each category throughout learning. We fit a linear regression model to each trajectory and extracted the best-fitting slopes. “Drifting” trajectories had significantly more negative slopes compared to “Stationary” trajectories (s = 71.0, p = 0.013, one-sided Mann-Whitney U-Test). On average, mice that exhibited stationary choice bias trajectories had category performance that increased symmetrically from their initial preference. As a result, their choice bias at the end of learning matched their initial choice bias (Figure 3C). Mice that had drifting choice bias trajectories, however, exhibited category performance that increased asymmetrically, often resulting in an eventual choice bias in the opposite direction from the initial choice bias (Figure 3D). The results of our clustering indicate that the extent to which the choice bias drifts throughout early learning is a key element in differentiating learning patterns.

**Figure 3.**
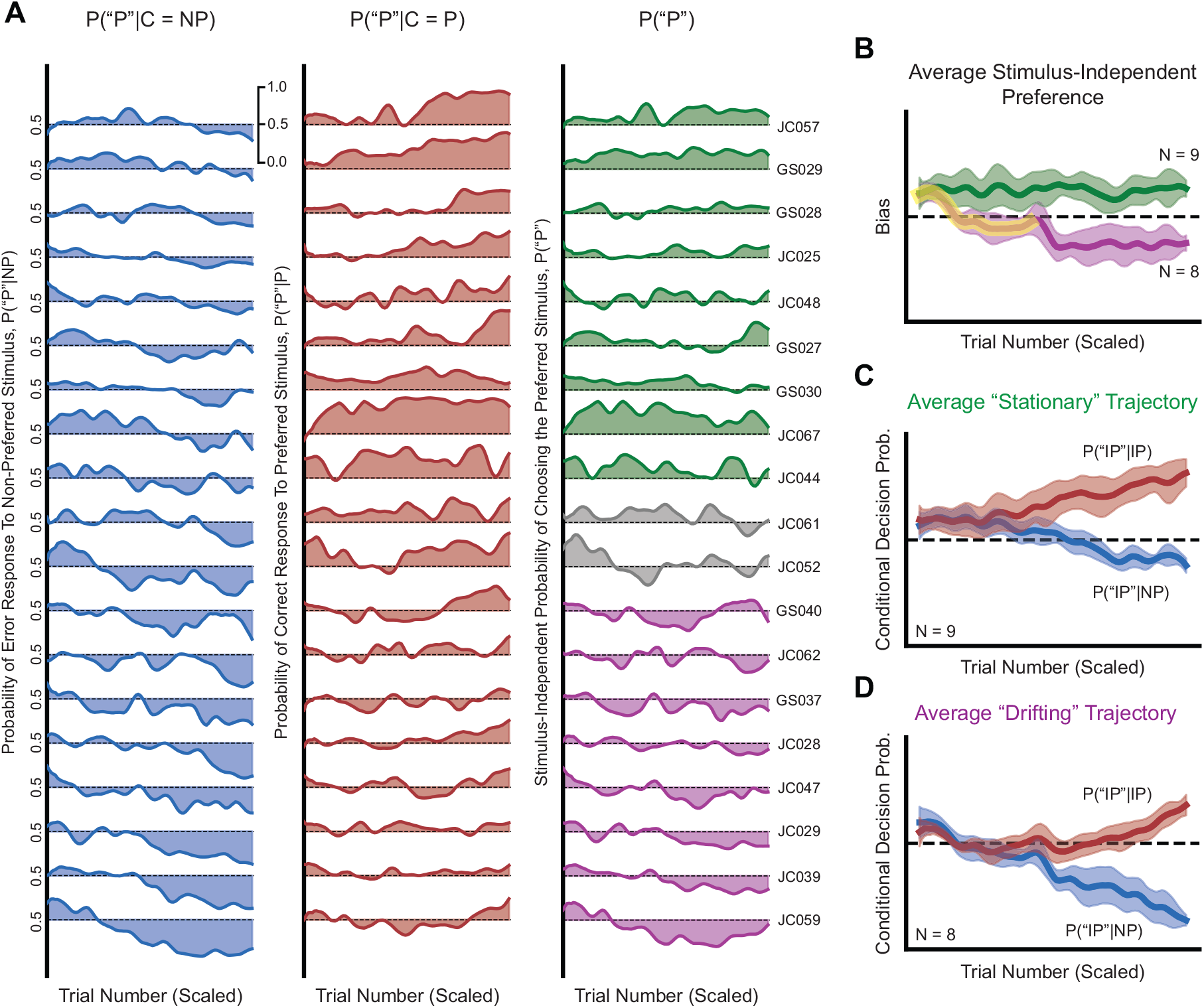
Early choice bias drift differentiates learning patterns. A, Left: Category-conditioned decision probabilities for the initially unpreferred category for each mouse. Center: Category-conditioned decision probabilities for initially preferred stimulus for each mouse. Right: Choice bias trajectory across learning for each mouse. Green curves indicate membership in “Stationary” cluster while purple coloring indicates membership in “Drifting” cluster. B, Average choice bias trajectory for each cluster. The average “Drifting” preference trajectory switches from the initially preferred category to the initially unpreferred category, whereas the average “Stationary” trajectory maintains the same category preference throughout learning. The drift away from the initial preference is highlighted in yellow. C, Average category-conditioned decision probabilities for mice labeled as having “Stationary” trajectories. On average, category performance increases symmetrically from initial preference. D, Average category-conditioned decision probabilities for mice labeled as having “Drifting” trajectories. On average, category performance increases asymmetrically, with a stimulus-independent shift in preference away from the initially preferred category. All trajectories were subsampled to 100 data points post-clustering for visualization. In all panels: error bars represent 95% CIs.

Next, we focused on what might drive individual variability in choice bias drift throughout learning. At the beginning of learning, subjects might not have a clear idea of the two stimulus categories. We hypothesized that the choice bias drift exhibited during early learning may be due to behavior that is independent of the presented stimulus. Because we trained our mice using reinforcement on every trial, we chose to use a reinforcement learning framework to probe this question (Figure 4A). The first model, which we refer to as the “choice-history” model, tested the extent to which the subjects update their associations based on their previous choice, independent of the presented stimulus. It has four free parameters: (1) a learning rate *α* that dictates how quickly the agent learns, (2) an overall bias *b* toward one of the two categories, (3) an initial bias *Q*_0_ toward one of the two categories, and (4) a choice-history parameter *β* (Figure 4A, Right panels). We also considered a null model in the same model family in which the choice-history parameter is removed (which we refer to as the “simple” model (Figure 4A, Left panels). Our hypothesis suggests that the choice-history model would be required to capture the behavior of mice that exhibit choice bias drift during early learning. Mouse trajectories that were labeled as not exhibiting choice bias drift would be better fit by the simple model. The fitting procedure maximizes the number of behavioral trials correctly predicted by the model (see Methods). To compare models from the same family but with different numbers of parameters, we compared the Bayesian Information Criterion (BIC) scores, which include a penalty for additional parameters. As we predicted, the behavioral performance of mice that were labeled as exhibiting choice bias drift during learning was more likely to be better fit by the choice-history model compared to mice not exhibiting choice bias drift (5/8 vs 0/9, p = 0.002, Prop Z-Test over BIC comparisons, Figure 4B).

**Figure 4.**
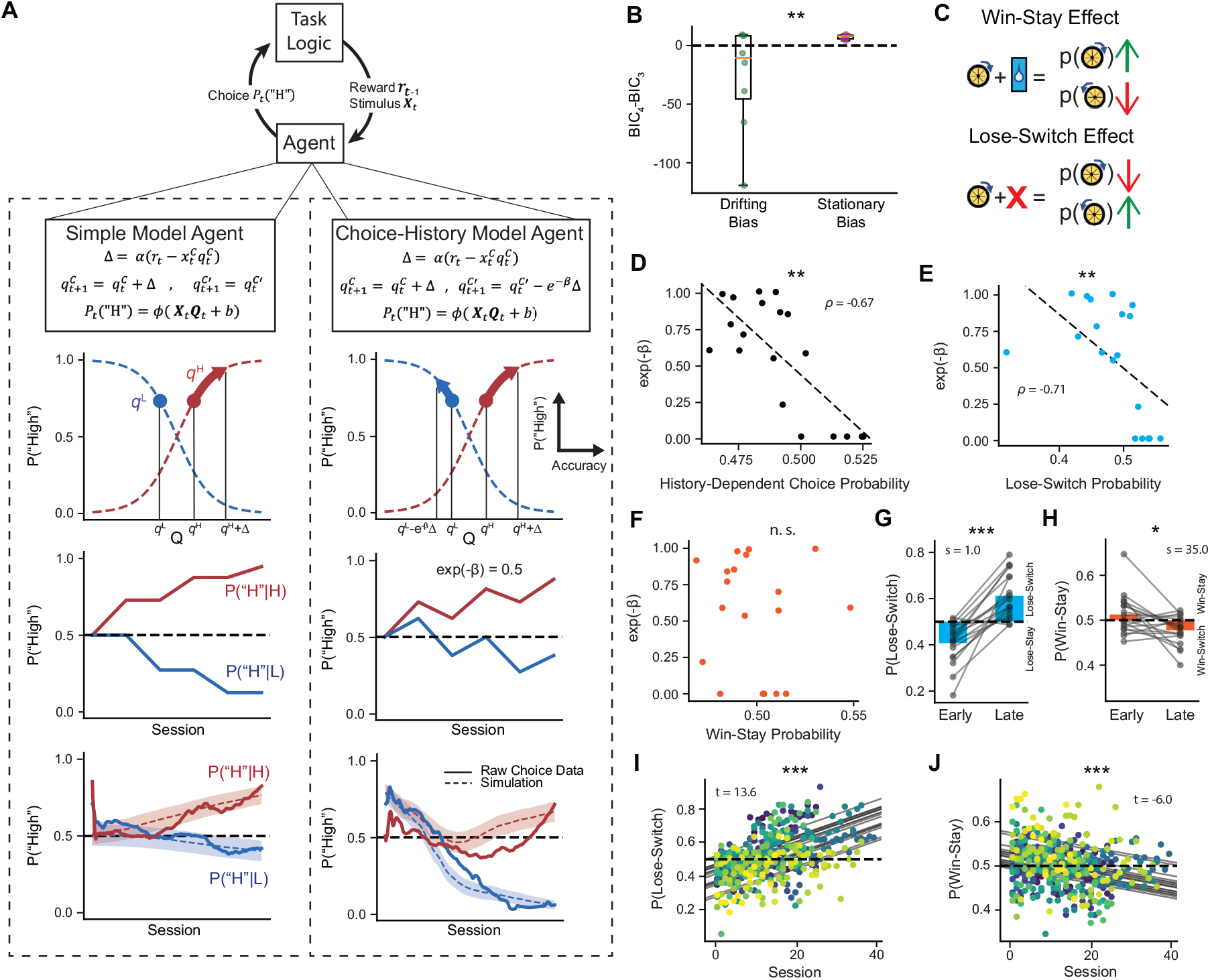
Group differences in choice bias drift during learning are due to subject-level variability in perseveration. A, Top: Schematic of reinforcement learning algorithm design. Second row: Equations governing agent behavior in response to stimuli for Simple Model (Left) and Choice-History Model with *e*^−*β*^ = 0.5 (Right). Third row: Example Q-Updating and logistic transformation in response to a correct “High” category trial. Left: For the Simple Model, only *q*^*H*^ is updated. Right: For the Choice-History Model, *q*^*L*^ is also updated, resulting in a slight bias towards the rewarded turn direction (“High”). Fourth row: Example category-conditioned probability updating for Simple Model (Left) and Choice-History Model with *e*^−*β*^ = 0.5 (Right) to the same series of stimuli and choices. A correlation between the category-conditioned probabilities emerges in the Choice-History Model. Fifth row: Model simulations based on parameters fit to example mouse better fit by Simple Model (Left) or Choice-History Model (Right). B, Simple model fits to trajectories that were clustered by the DTW as having “drifting choice bias” were better improved by the addition of the choice-history parameters compared to trajectories that were clustered by the DTW as having “non-drifting choice bias”. Improvement was quantified by comparing the Bayesian Information Criterion (BIC) of the two fits. A lower BIC indicates a better model fit. C, Diagram explaining Win-Stay and Lose-Switch effects. D, Choice-history parameter captured inter-subject variability in choice-history. E, Choice-history parameter captured inter-subject variability in tendency to switch after an incorrect trial. F, Choice-history parameter was not correlated with behavior following a correct trial. G, Mice tended to exhibit more lose-switch behavior during their final 3 training sessions compared to their first three training sessions. H, Mice tended to exhibit less win-stay behavior during their final 3 training sessions compared to their first three training sessions. I, Lose-switch behavior increases as a function of training session number. J, Win-switch behavior decreases as a function of training session number. In I and J: Grey lines indicate individual trend lines from OLS model with different intercepts across mice. Colors indicate specific mouse, N = 19 mice. In all plots: n.s., not significant; *p<0.05; **p<0.01; ***p<0.001; ****p<0.0001.

To determine what elements of choice history the choice-history model captures, we calculated each mouse’s average probability of responding according to the previous choice and reward (either “staying” after a win (win-stay), or “switching” after a loss (lose-switch), Figure 4C). This value is strongly negatively correlated with the choice-history parameter (*ρ* = -0.67, p = 0.002, Spearman correlation), indicating that the choice-history parameter captured switching after a win (win-switch) or staying after a loss (lose-stay) (Figure 4D). We found that the choice-history parameter was correlated with lose-stay behavior (*ρ* = -0.71, p = 0.001, Spearman correlation), but not win-switch (*ρ* = -0.07, p = 0.78, Spearman correlation, Figure 4E-F).

Lose-stay behavior is a form of perseverative behavior^21^. Perseverative behavior refers to a choice bias toward repeating the previous decision. Whereas win-stay behavior is also perseverative as the choices are repeated, our choice-history model specifically captures perseveration in response to incorrect trials (Figure 4E-F). Analyzing the extent of choice-history correlations across sessions revealed more perseverative behavior following incorrect trials in early sessions (first three sessions) compared to late sessions (final three sessions, s = 1, p = 7.6e-06, Wilcoxon Sign-Rank) and more win-stay behavior in early sessions compared to late sessions (s = 35, p = 0.014, Wilcoxon Sign-Rank, Figure 4G-H). This high rate of perseveration following incorrect trials during early learning matched the region of learning where the choice bias drift occurs (Figure 3B). In addition, we found that the perseverative behavior following incorrect trials increased with the number of sessions completed (t = 13.6, p = 4.0e-35, OLS model, Figure 4I), while the win-stay behavior decreased with the number of sessions completed (t = -6.0, p = 5.2e-09, OLS model, Figure 4J). Our results support the hypothesis that the group differences in choice bias drift during learning are due to subject-level variability in attention to choice history, specifically perseverative tendencies to repeat the losing choice following incorrect trials.

## Discussion

We examined the relationship between the learning trajectories of mice for an auditory categorization task and resulting categorization parameters. This task was designed to encourage stimulus abstraction by exposing subjects to categories consisting of a range of tone burst stimuli (Figure 1). Different mice exhibited different patterns in their learning trajectories (Figure 3). Mice varied in their choice bias before they were introduced to the category-ambiguous stimuli (Figure 3). The mouse behavior could be interpreted as a combination of reward-driven learning and choice bias (Figure 2). We found that inter-individual variability in this choice bias was predictive of the stability of the category boundary location (Figure 2), suggesting that the learning dynamics contribute to learned category boundaries. Next, to study individual variability in behavioral dynamics during learning, we applied time-series clustering to the learning trajectories of mice. Our results identified across-learning drift in bias as a prominent pattern in learning dynamics (Figure 3). This suggests that that both rewards and choice biases shape the learning of categories and influence the resulting category boundaries. Furthermore, this drift in bias and individual response variability depended on the history of the mouse’s previous responses, specifically the tendency to perseverate following incorrect decisions (Figure 4). These results indicate that long-term trends in choice bias during category learning reflect a preference that the mice rely on when drawing category boundaries.

Individuals often learn and build category rules through trial and error, using a variety of strategies to ensure that incoming stimuli are assigned their proper labels. Some of these strategies, such as basing responses on previous choices or previous reward contingencies, are at least partially driven by factors independent from the stimuli presented. Although recent work has extracted behavioral significance from stimulus-independent dynamics within sessions in trained rodents^10^, it was previously unknown whether there was a behavioral significance to the long-term trajectories of this stimulus-independent component. Here, we find that the stimulus-independent components of learning trajectories inform resulting category boundaries. In addition, whereas it has been demonstrated that elements of behavior during early learning are predictive of future performance on the same stimuli^5^, we additionally find that individual variability in behavioral dynamics during learning informs the categorization of novel category-ambiguous stimuli.

An important implication of this study is that the preference that drives categorization behavior to category-ambiguous stimuli can be tracked throughout learning. In real-world scenarios, subjects learn to categorize not just for the sake of being rewarded for correct categorization to known stimuli but to more efficiently understand and interact with their environment^22^. One caveat in studying rodent behavior is that we cannot directly ask them to assign labels, rather we train them to respond to stimuli with distinct movements. In our task, we trained them to turn the wheel clockwise and counterclockwise. This is not unlike a table tennis player learning to use a forehand or backhand stroke. As a table tennis player improves their forehand and backhand strokes, their behavior in response to shots in front of their body changes and they exhibit individual variability in stroke preference. A subject skilled in both strokes but who considers their backhand a more consistent option might exhibit a preference towards using that stroke in response to shots with ambiguous trajectories. A player who originally preferred their backhand but, through training, built up confidence in their forehand, might exhibit a preference towards using their forehand in response to ambiguous shots because of that recently acquired confidence. Here, we established a method for tracking the development of category preferences for individual subjects as they interact with and learn categories.

We found that choice bias can be traced throughout learning and is predictive of categorization behavior. The long-term trend in this bias is a notable pattern across mice which our model attributes to initial bias and weighting of previous choices. However, variability driving this initial bias (such as handedness, position, and strategies picked up during early habituation) are beyond the scope of this work. In addition, although our model identified perseverative behavior as involved in the long-term trend, recent work found dynamic bias at a shorter time-scale and in trained animals^10,11^ as linked to an exploration vs. exploitation process. The relationship between choice bias during learning and category boundaries that we found is consistent with the interpretation of exploration being reflected in the dynamic choice bias. As both choice-history weighting and exploration would be able to account for the dynamics we encountered, future work should explore designing a new task condition where reward effects are asymmetrically manipulated during training to determine whether the choice bias is driven by reward effects.

Humans learning novel categories with feedback tend to repeatedly probe the same category as opposed to interleaving queries about multiple categories, and predict higher future performance when practicing in this way^6^. Similarly, mice that exhibit long-term drift in choice bias may be alternating periods of “studying” the outcomes of the two response options: clockwise and counterclockwise wheel turns. during early learning, this repeated study would result in perseveration, the tendency to repeat previous choices. We find that win-stay behavior tends to decrease in prevalence throughout learning, while lose-switch behavior increases in prevalence. Both effects can be explained by perseveration decreasing throughout learning. Perseveration is a fundamental aspect of human reinforcement learning^23^ and rodent category learning^8^. Our finding that perseveration is strongest early in category learning and decreases throughout training matches previous work with rodents^8^. Due to the inclusion of correction trials in our task design, the emergence of lose-switch behavior throughout learning likely arises in part from a decrease in perseveration (as indicated by the decrease in win-stay behavior) and in part from the mice learning the task structure. Future work may consider a training regime without correction trials to allow for a detailed analysis of these contributions to lose-switch behavior.

Whereas the general decrease in win-stay behavior throughout learning is consistent with previous results^8^, our finding that most of the mice exhibited consistent win-switch behavior late during learning was unexpected. It was unexpected because win-switch is not a heuristic traditionally associated with sequential learning, unlike win-stay and lose-switch^24^. It is possible that this switching behavior was a consequence of the high uncertainty during early learning, which encouraged switching because of the high probability of incorrect choices. A bias toward switching following a correct choice would also be consistent with a tendency to explore multiple options. Future experiments could test explicitly whether mice can learn to favor exploration over consolidation of learned responses by varying reward contingencies across different stages of the task.

The wheel-based 2AFC task we implemented is a relatively difficult task for mice to learn, resulting in high inter-subject variability in the time taken to complete training (ranging from 4000 - 15000 trials). The computational methods in this study are invariant to the actual number of trials taken for each mouse to learn the task. Although the wheel-based 2AFC task offers several advantages over other 2AFC tasks, such as the separation of choice action (turning the wheel) from reward gathering action (licking at the lick port) and the continuous readout from the wheel^12^, other 2AFC tasks, such as dual-lick port setups, offer shorter and more consistent learning times^25^. Future work may consider the effect of task difficulty on inter-subject variability during learning dynamics.

Our task paradigm and modeling approach allowed us to identify patterns in category preferences as they evolved throughout the learning process. These patterns primarily differed based on whether the direction of the choice bias was consistent or switched throughout training. Our finding that auditory cortex inactivation impaired categorization behavior raises the question of whether the auditory cortex maintains a representation of the category boundary throughout category learning. Future work might use our methodology combined with long-term electrophysiological recordings to determine how category encoding in the sensory cortex develops over learning.

## Methods

### Animals

Experiments were performed in adult male (n=8) and female (n=11) mice. Most of the mice (n = 17) were B6.CAST-Cdh23Ahl+ (Stock No. 002756) mice (The Jackson Laboratory; age 12–15 weeks; weight 20–30g). Two non-Cdh23 mice were included (1 male, 1 female), their removal does not change the results. Some of the mice were crossed with other cell-type specific cre lines, as detailed in Table 2. All mice were housed with 2-5 mice per cage, at 28°C on a 12-h light-dark cycle with food and water provided ad-libitum pre-surgery, and water restricted post-surgery as described below. All experiments were performed during the animals’ dark cycle. All experimental procedures followed NIH guidelines and were approved by the Institutional Animal Care and Use Committee at the University of Pennsylvania.

**Table 1.**
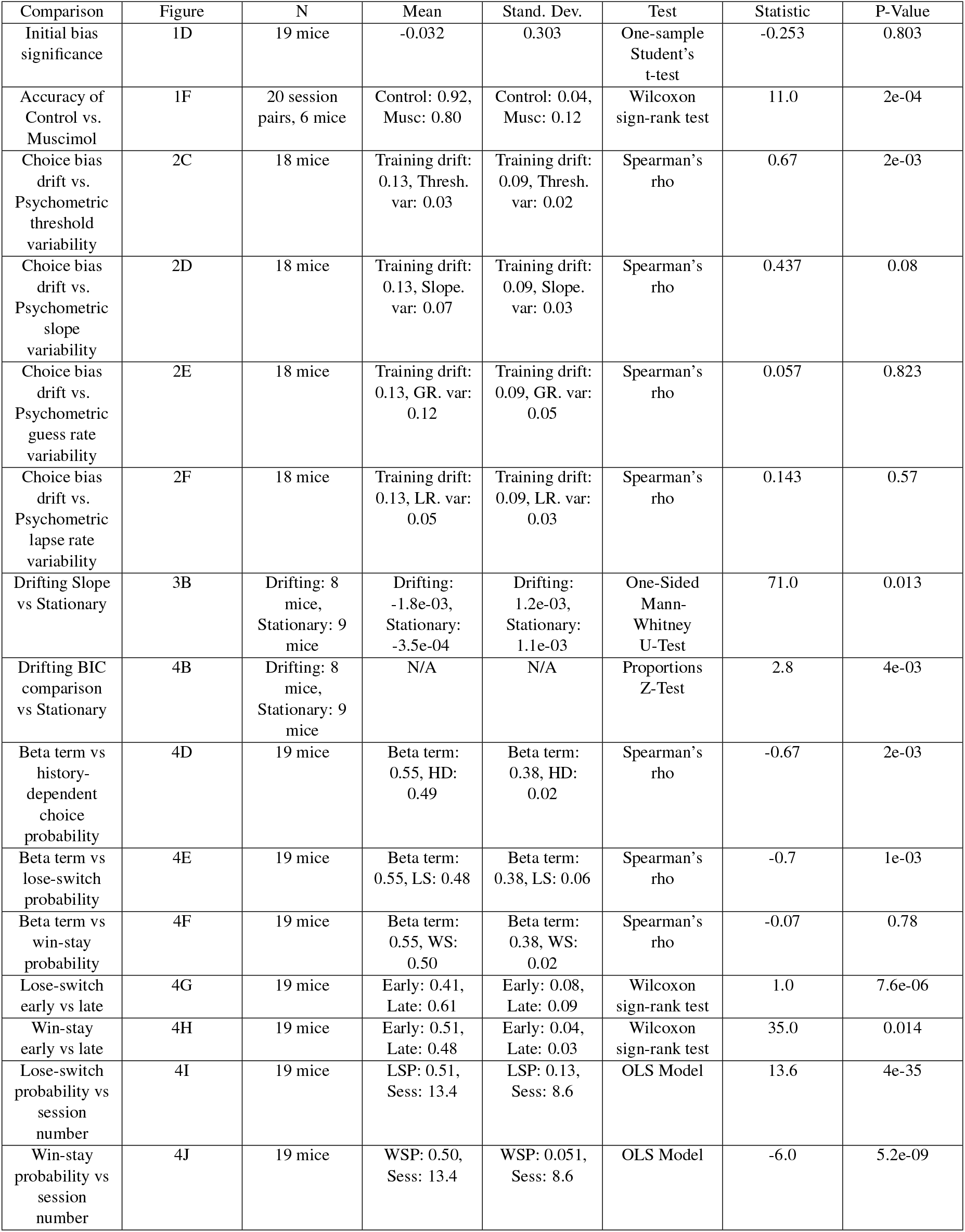
Statistical analyses.

**Table 2.** Mouse strains and sex.

| Strain | N [Female] | N [Male] |
| --- | --- | --- |
| Cdh23 | 8 | 6 |
| Cdh23 x CAMKII-cre | 1 | 1 |
| Cdh23 x VGAT | 1 | 0 |
| Rosa x VGAT | 1 | 1 |
| Total | 11 | 8 |

### Surgery

Mice were anesthetized using 1-3% isoflurane. All mice were administered subcutaneous doses of buprenorphine (Buprenex, 0.05–0.1 mg/kg) as analgesia, dexamethasone (0.2 mg/kg) to reduce brain swelling, and bupivacaine (2 mg/kg) as local anesthesia. Mice were implanted with 1-2 skull screws in the cerebellum, and a headplate was mounted on the skull as previously described^26^. An antibiotic (Baytril, 5mg/kg) and analgesic (Meloxicam, 5 mg/kg) were administered daily (for 3 days) during recovery. For mice undergoing muscimol injections, infusion cannulas (26 GA, PlasticsOne, C315GMN-SPC mini) were implanted bilaterally over the auditory cortices at a location of -2.6mm posterior to Bregma/4.3 mm lateral and a depth of 500 microns. Dummy cannulas (PlasticsOne, C315DCMN-SPC mini with 0.5mm projection depth) were partially screwed into the guide cannulas to minimize cannula blockage in the time between surgery and the injection sessions.

### Auditory Cortex Inactivation

Prior to injection sessions, two Hamilton syringes (10uL Gaslight #1701) and tubing (C313CT 023×050 PE50) were backfilled with mineral oil. We attached a sterilized infusion cannula to the end of each tube and drew up either 500 nL of either muscimol (diluted with 1x PBS to 1.5 mg/mL, which was used on inactivation sessions) or 1x PBS (used on control sessions)^27^. We headfixed the mouse, removed the dummy cannulas, and then inserted the loaded infusion cannulas into the guide cannulas. We infused 450 nL of the solution bilaterally at a rate of 250 nL/minute. We then removed the infusion cannulas and re-inserted the sterile dummy cannulas. The mice then rested in their home cage for 45 minutes before their behavioral session. Control sessions were followed by injection sessions, with data collected from 20 session pairs across 6 mice. The number of mice was determined using a power analysis in MATLAB.

### Water Restriction

To establish a baseline weight, each mouse was monitored for 3 days post-operation. Once the mice regained weight, water deprivation began with a daily ration of 120 µL/g, which gradually decreased to 40–50 µL/g. During the task, if mice did not receive their full ration, the remainder of their ration was provided in their home cage. Mouse weight, relative to baseline, was monitored throughout water restriction. A health score was also calculated every day, and mice were assigned a treatment plan if this score dropped or if the mouse lost more than 20% of their baseline weight^25^.

### Behavioral Apparatus

During the 2AFC task, the mouse was head-fixed in a custom-built, acoustically isolated chamber. A wheel (12-mm Hub Wheels, GDOOL) attached to a rotary encoder (COM-10932 ROHS, Sparkfun) monitored wheel activity, whereas a capacitive touch sensor (AT42QT1010, SparkFun) soldered to a lick spout monitored lick activity. Water rewards were dispensed from a gravity-fed reservoir calibrated to deliver approximately 4-5µL of water per reward. An Arduino Uno microprocessor was used for monitoring of the rotary encoder, reward delivery, and task logic. Custom MATLAB software communicating with the Arduino controlled the sampling of categories and tone burst frequencies, stimulus generation, and online data collection and analysis. For the stimulus generation, a sound card (Lynx E44, Lynx Studio Technology, Inc.) or a National Instruments card (NI PCIe-6353) was used to convert the MATLAB-generated acoustic waveforms into analog signals. An ultrasonic transducer (MCPCT-G5100-4139, Multicomp) delivered the analog signals. A 1/4-inch condenser microphone (Brüel & Kjær), positioned at the estimated location of the mouse’s ear, was used to calibrate the transducer to have a flat frequency response between 3 kHz and 80 kHz^28,29^.

All stimuli were sampled at 192 kHz or 200 kHz and 32-bit resolution. A set of tone-burst stimuli was generated at different frequencies. In some experiments (n = 87 training sessions), 60 frequencies were sampled, logarithmically spaced from 8 to 32 kHz. For these experiments, “low-frequency stimuli” were defined as frequencies between 8 and 10.8 kHz, “probe stimuli” were defined as frequencies between 10.8 and 23 kHz, and “high-frequency stimuli” were defined as frequencies between 23 and 32 kHz. In the remaining experiments (n = 354 training sessions), 30 frequencies were sampled, logarithmically spaced from 6 to 28 kHz. For these experiments, “low-frequency stimuli” were defined as frequencies between 6 - 10 kHz, “probe stimuli” were defined as frequencies between 10 and 17 kHz, and “high-frequency stimuli” were defined as frequencies between 17 - 28 kHz. Effects were not dependent on the stimulus set.

### Behavioral Training

Each mouse underwent three stages of behavioral training: 1) water restriction and habituation, 2) 2AFC training, and 3) psychometric testing. During water restriction, mice were habituated to head-fixation in the behavioral chambers and received water through the lick spout, getting a drop of water for licks separated by more than 2 seconds. During this stage, the wheel was taped in place. After the mouse began to receive its entire ration of 1-2 mL of water through the task rewards, a second habituation stage was initiated (typically after 2-3 days). In this second habituation stage, we trained mice to turn the wheel 30° in alternating directions and to hold the wheel still for 500 ms between turns. The mouse was rewarded with a drop of water for each turn. Moving the wheel during the holding period restarted the hold time. When the mouse began to receive its entire ration of 1-2 mL of water through the task rewards, the 2AFC training stage was initiated (this stage typically took 5-8 days). In this training stage, mice were instructed to turn the wheel in response to a 500 ms tone burst. The mice were equally likely to be assigned to associate clockwise turns with low-frequency (6-10 kHz) or high-frequency (17-28 kHz) stimuli. Mice had 3 s from stimulus onset to register a wheel turn. An incorrect turn resulted in a noise burst and a 6-s time-out period, whereas a “no response” trial resulted in the previous trial being repeated. Mice were instructed to hold the wheel still for a period randomly drawn between 1 and 2 s in order for the next trial to commence. The mice were made aware of the next trial beginning upon hearing the auditory stimulus. A single session consisted of repeated trials until the mouse either began to stop responding or received its full water ration (200-300 trials). We monitored behavioral performance during training, and considered mice as trained after they completed two consecutive sessions with >75% accuracy for each category. For analysis purposes, only the first of these two sessions was included in the learning trajectories. After the mice completed two sessions with accuracies to each category above-threshold, they were moved to the psychometric testing stage. In this testing stage, we introduced novel tone burst stimuli with center frequencies in between the two frequency ranges used for the training stage (10-17 kHz) to study how the mice draw category boundaries between the two categories. These novel stimuli were presented on 20% of trials.

### Data Analysis

#### Extracting Stimulus-Independent Dynamics

For all training trials, the stimuli were separated based on their category. Tone bursts from the high-frequency category were transformed into “evidence towards high” through normalization based on all tone bursts experienced by the mouse during that session. For example, the tone burst with the highest frequency would have an “evidence towards high” of 1, while the lowest tone burst drawn from the high distribution might have an “evidence towards high” of 0.66. The same was done for trials with tone bursts drawn from the low-frequency category. These values were then transformed using a tau distribution with *τ* = 2 to reflect the fact that mice generally show stability within the trained stimulus region. PsyTrack, a method for computing the trajectories of model parameters, was used to fit a generalized linear model to this behavioral data using three dynamic variables-two category weights and a dynamic choice bias^8^.

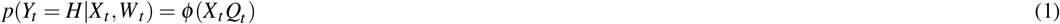

The weights *W* evolve gradually over time, with each weight *w*_*i*_ having an individually fit rate of change.

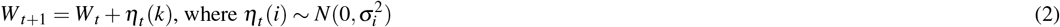

#### Quantifying Categorization Behavior

To quantify the categorization behavior of the mice, we fit the testing data with a 4-parameter psychometric function. Each decision is modeled as either being the result of a decision function characterized by the parameters *µ* (threshold) and *σ* (slope) (with probability 1 − *γ*) or, with probability *γ*, the result of lapse behavior, where the decision is “High” with probability *λ* and “Low” with probability 1 − *λ*.

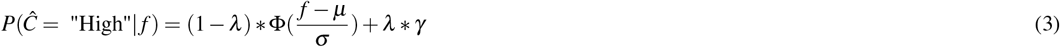

We used the PyBADS toolbox to fit this model to behavioral data from individual testing sessions for each mouse^30^. To determine whether the dynamic bias informed categorization under uncertainty, we calculated the variability of the GLM choice bias over the final day of training for each mouse and determined the strength and direction of the relationship between this value and the variability of the category threshold over the first 5 testing sessions using the Spearman correlation. These analyses were performed using the SciKit-Learn package^31^.

#### Generating Learning Trajectories

We generated category-conditioned accuracy traces by first separating out mouse responses to high-frequency and low-frequency trials. These traces consisted of responses for sessions where the mice were exposed to the training stimuli, up to and including the first day that the accuracy for each category passed 75%. These category-conditioned traces were smoothed by taking a rolling average and then subsampled every 50 trials. We then calculated the most chosen category for each mouse over their first session and defined that as the mouse’s “preferred category”.

#### Clustering

We first calculated the probability that a mouse correctly responded to low-category stimuli *P*(*H*|*L*) and high-category stimuli *P*(*H*|*L*). We applied a smoothing algorithm over 50 trials to generate estimated category-conditioned accuracy traces. To generate the trace estimating the probability of the mouse responding “High”, we averaged the two category-conditioned accuracy traces.

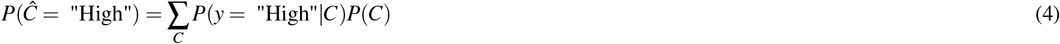

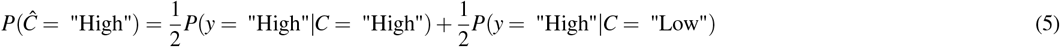

To remove clusters that differed based on their direction of stimulus-independent dynamics, we assigned a direction to the vectors such that the positive direction was towards their initial bias over their first training session. The resulting vectors, *P*(*Ĉ* = “Preferred Category”), were clustered using Dynamic Time Warping (DTW) classification. It is preferable to Principle Component Analysis in situations in which the data being clustered is smoothed over time^32^. We utilized the Python package Tslearn for the DTW clustering^33^. We clustered the vectors 100 times and calculated the proportion of the runs that each pair of mice were clustered together. We used 70% co-clustering as the threshold for a pair being consistently clustered together and found that 17/19 mice were consistently clustered into the same groups, while 2 mice co-clustered with each other but not with any other mice. These 2 mice were not considered in analyses based on clustering.

#### Learning Model

We constructed a simple reinforcement learning model to capture the learning dynamics of the mice. For this model, stimuli were binarized as “low” or “high” based on their category membership. Stimulus evidence on trial *t* was encoded as *X*_*t*_ = [*x*_*L*_, *x*_*H*_], where *x*_*L*_ is the evidence towards low in a given trial, and *x*_*H*_ is the evidence towards high. Low-frequency stimuli were encoded as *X*_*t*_ = [−1, 0], whereas high-frequency stimuli were encoded as *X*_*t*_ = [0, 1]. To simulate the subject’s choice, the stimulus evidence towards the high category *E*_*t*_ was multiplied by the subject’s stimulus-action associations *Q*_*t*_ = [*q*_*L*_, *q*_*H*_], and shifted by a constant bias term *b*. To reflect that the bias is not necessarily static, we allowed for an initial stimulus action-association bias, *Q*_init_, that varied across subjects.

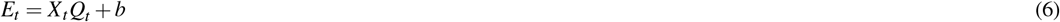

The probability that the agent chooses the high category based on the stimulus evidence 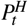 was calculated by transforming the stimulus evidence into a probability space using the logistic transformation.

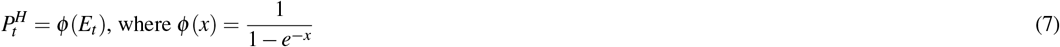

We fit this model by calculating the log-likelihood *LL* for each trial. *LL* was calculated by comparing the simulated agent’s decision probability 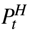 with the actual decision of the mouse on the given trial. The reward state for the trial, *r*_*t*_, was determined by the true decision of the mouse, not the simulated decision probability. This reward state was used to update the stimulus-action association 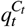 using a delta-rule. *C*_*t*_ encodes the stimulus category presented on trial *t*, and *α* represents the learning rate.

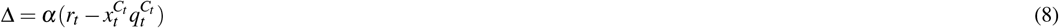

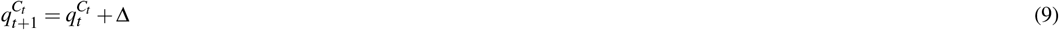

We used this simple model as a baseline for model comparison. In this model, the three free parameters are the overall preference *b*, the initial bias *Q*_init_, and the learning rate *α*. We chose these three parameters because they control the shape of the learning trajectory. To capture the stimulus-independent dynamics during early learning, we modified this simple model by adding a choice-history coefficient, *β*. This parameter controlled the extent to which the updating associated with the reward state caused the agent to change their overall valuation of the previous action, independent of the presented stimulus category. This additional parameter allows for more complex stimulus-independent trajectories during early learning.

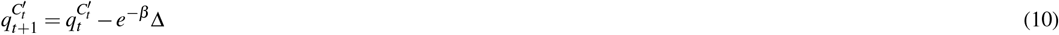

Here, 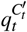 is the stimulus-action association of the non-presented category. The negative updating results in a shift towards the same turn direction as the stimulus-action association of the presented category. For large *e*^−*β*^ terms, 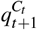 and 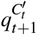 are both shifted towards the same turn direction with similar magnitudes, meaning little category learning has occurred.

#### Choice-History Effects

For each mouse, across the training regime, we calculated the probability of turning in the previous trial’s “correct” direction (win-stay following a reward or lose-switch following an error trial). We additionally calculated the probability of the mouse exhibiting win-stay and lose-switch behavior, separately. We determined the strength and direction of the correlation between these probabilities and the *β* term using the Spearman coefficient. We calculated the individual win-stay and lose-switch probabilities for each mouse for each session and compared the first 3 and final 3 training sessions for each mouse using a Wilcoxon Sign-Rank test. These analyses were performed using the SciKit-Learn package^31^. For calculating the main effects of session number on win-stay and lose-switch probabilities, we used an Ordinary Least Squares (OLS) model with different intercepts for each mouse. There was a high level of variability in the initial level of win-stay and lose-switch probabilities across mice, and this model allowed us to determine whether there was an overall effect of session number while acknowledging this variability. We used the p-values associated with whether the slope was different from zero to quantify whether there was a main effect for each analysis. These analyses were performed using the statsmodels package^34^.

## Data and Code Availability

The data and code associated with this paper are publicly available. The code can be accessed at https://github.com/geffenlab/CategorizationStrategies, while the data has been made available on Dryad at https://doi.org/10.5061/dryad.73n5tb359.

## Acknowledgments

This research is supported by the grants to M.N.G., K.P.K., and Y.E.C. (NIH NINDS R01NS113241), grants to M.N.G. (NIH NIDCD R01DC015527 and R01DC014479) and fellowship to J.S.C. (CANAC T32DC016903-01). We would like to thank members of the Geffen laboratory for their feedback.

## Author Contributions

J.S.C., C.F.A., K.C.W., J.S., K.P.K, Y.E.C. and M.N.G. conceived the experiments. J.S.C., M.X. and G.E. conducted the experiments. J.S.C. analyzed the results. J.S.C., K.P.K., Y.E.C. and M.N.G. wrote the paper.

## Guidelines

The authors confirm that the study is reported in accordance with ARRIVE guidelines (https://arriveguidelines.org).

## References

1. Tsunada, J. & Cohen, Y. E. Neural mechanisms of auditory categorization: From across brain areas to within local microcircuits, DOI: 10.3389/fnins.2014.00161 (2014).

2. Freedman, D. J., Riesenhuber, M., Poggio, T. & Miller, E. K. Categorical representation of visual stimuli in the primate prefrontal cortex. Science 291, 312–316, DOI: 10.1126/science.291.5502.312 (2001).

3. Gifford, A. M., Cohen, Y. E. & Stocker, A. A. Characterizing the impact of category uncertainty on human auditory categorization behavior. PLoS computational biology 10, e1003715, DOI: 10.1371/journal.pcbi.1003715 (2014).

4. Kluender, K. R., Coady, J. A. & Kiefte, M. Sensitivity to change in perception of speech. Speech Commun. 41, 59–69, DOI: 10.1016/S0167-6393(02)00093-6 (2003).

5. Liebana Garcia, S. et al. Striatal Dopamine Reflects Long-term Learning Trajectories. bioRxiv DOI: 10.32470/ccn.2023.1477-0 (2023).

6. Bjork, R. A., Dunlosky, J. & Kornell, N. Self-regulated learning: Beliefs, techniques, and illusions. Annu. Rev. Psychol. 64, 417–444, DOI: 10.1146/annurev-psych-113011-143823 (2013).

7. Hélie, S., Shamloo, F. & Ell, S. W. The effect of training methodology on knowledge representation in categorization. PLoS ONE 12, 1–23, DOI: 10.1371/journal.pone.0183904 (2017).

8. Roy, N. A., Bak, J. H., Akrami, A., Brody, C. D. & Pillow, J. W. Extracting the dynamics of behavior in sensory decision-making experiments. Neuron 109, 597–610, DOI: 10.1016/J.NEURON.2020.12.004 (2021).

9. Ashwood, Z. C. et al. Mice alternate between discrete strategies during perceptual decision-making. Nat. Neurosci. 25, 201–212, DOI: 10.1038/s41593-021-01007-z (2022).

10. Pisupati, S., Chartarifsky-Lynn, L., Khanal, A. & Churchland, A. K. Lapses in perceptual decisions reflect exploration. eLife 10, 1–27, DOI: 10.7554/ELIFE.55490 (2021).

11. Zhu, Z. & Kuchibhotla, K. V. Performance errors during rodent learning reflect a dynamic choice strategy. Curr. Biol. 34, 2107–2117, DOI: 10.1016/j.cub.2024.04.017 (2024).

12. Burgess, C. P. et al. High-Yield Methods for Accurate Two-Alternative Visual Psychophysics in Head-Fixed Mice. Cell Reports 20, 2513–2524, DOI: 10.1016/j.celrep.2017.08.047 (2017).

13. King, A. J., Teki, S. & Willmore, B. D. Recent advances in understanding the auditory cortex., DOI: 10.12688/F1000RESEARCH.15580.1 (2018).

14. Krall, R. F. et al. Primary auditory cortex is necessary for the acquisition and expression of categorical behavior. bioRxiv 2024.02.02.578700 (2024).

15. Yin, P., Strait, D. L., Radtke-Schuller, S., Fritz, J. B. & Shamma, S. A. Dynamics and Hierarchical Encoding of Non-compact Acoustic Categories in Auditory and Frontal Cortex. Curr. Biol. 30, 1649–1663, DOI: 10.1016/j.cub.2020.02.047 (2020).

16. Tsunada, J., Liu, A. S., Gold, J. I. & Cohen, Y. E. Causal contribution of primate auditory cortex to auditory perceptual decision-making. Nat. neuroscience 19, 135, DOI: 10.1038/NN.4195 (2016).

17. Xin, Y. et al. Sensory-to-Category Transformation via Dynamic Reorganization of Ensemble Structures in Mouse Auditory Cortex. Neuron 103, 909–921, DOI: 10.1016/j.neuron.2019.06.004 (2019).

18. Selezneva, E., Scheich, H. & Brosch, M. Dual Time Scales for Categorical Decision Making in Auditory Cortex. Curr. Biol. 16, 2428–2433, DOI: 10.1016/j.cub.2006.10.027 (2006).

19. Niwa, M., Johnson, J. S., O’Connor, K. N. & Sutter, M. L. Differences between primary auditory cortex and auditory belt related to encoding and choice for AM sounds. J. Neurosci. 33, 8378–8395, DOI: 10.1523/JNEUROSCI.2672-12.2013 (2013).

20. Bizley, J. K. & Cohen, Y. E. The what, where and how of auditory-object perception. Nat Rev Neurosci 14, 693–707, DOI: 10.1038/nrn3565 (2013).

21. Fuchs, B. A. et al. Decision-Making Processes Related to Perseveration Are Indirectly Associated With Weight Status in Children Through Laboratory-Assessed Energy Intake. Front. Psychol. 12, 1–18, DOI: 10.3389/fpsyg.2021.652595 (2021).

22. Markman, A. B. & Ross, B. H. Category Use and Category Learning. Psychol. Bull. 129, 592–613, DOI: 10.1037/0033-2909.129.4.592 (2003).

23. Sugawara, M. & Katahira, K. Dissociation between asymmetric value updating and perseverance in human reinforcement learning. Sci. Reports 11, 1–13, DOI: 10.1038/s41598-020-80593-7 (2021).

24. Goodnow, J. J. & Pettigrew, T. F. Effect of prior patterns of experience upon strategies and learning sets. J. Exp. Psychol. 49, 381–389, DOI: 10.1037/h0049350 (1955).

25. Guo, Z. V. et al. Procedures for behavioral experiments in head-fixed mice. PLoS ONE DOI: 10.1371/journal.pone.0088678 (2014).

26. Angeloni, C. F. et al. Cortical efficient coding dynamics shape behavioral performance. bioRxiv 2021.08.11.455845, DOI: 10.1101/2021.08.11.455845 (2021).

27. Misane, I., Kruis, A., Pieneman, A. W., Ögren, S. O. & Stiedl, O. GABA A receptor activation in the CA1 area of the dorsal hippocampus impairs consolidation of conditioned contextual fear in C57BL/6J mice. Behav. Brain Res. 238, 160–169, DOI: 10.1016/j.bbr.2012.10.027 (2013).

28. Carruthers, I. M., Natan, R. G. & Geffen, M. N. Encoding of ultrasonic vocalizations in the auditory cortex. J. Neurophysiol. 109, 1912–1927, DOI: 10.1152/jn.00483.2012 (2013).

29. Carruthers, I. M. et al. Emergence of invariant representation of vocalizations in the auditory cortex. J. Neurophysiol. 114, 2726–2740, DOI: 10.1152/jn.00095.2015 (2015).

30. Singh, G. S. & Acerbi, L. PyBADS: Fast and robust black-box optimization in Python. J. Open Source Softw. 9, 5694, DOI: 10.21105/joss.05694 (2024).

31. Pedregosa, F. et al. Scikit-learn: Machine Learning in Python. J. Mach. Learn. Res. 12, 2825–2830 (2011).

32. Shinn, M. Phantom oscillations in principal component analysis. Proc. Natl. Acad. Sci. United States Am. 120, 1–11, DOI: 10.1073/pnas.2311420120 (2023).

33. Tavenard, R. et al. Tslearn, A Machine Learning Toolkit for Time Series Data. J. Mach. Learn. Res. 21, 1–6 (2020).

34. Seabold, S. & Perktold, J. Statsmodels: Econometric and Statistical Modeling with Python. SciPy DOI: 10.25080/MAJORA-92BF1922-011 (2010).

